# A highly sensitive LC-MS/MS method for quantitative determination of 7 vitamin D metabolites in mouse brain tissue

**DOI:** 10.1101/2022.02.21.481384

**Authors:** Andrea Stephenson, Ben Hunter, Paul Nicholas Shaw, Nur Sofiah Abu Kassim, Rob Trengrove, Ryu Takechi, Virginie Lam, John Mamo

**Author notes:** **Correspondence:** John Mamo.

## Abstract

Despite its critical role in neurodevelopment and brain function, vitamin-D (vit-D) homeostasis, metabolism and kinetics within the central nervous system remain largely undetermined. Thus, it is of critical importance to establish an accurate, highly sensitive and reproducible method to quantitate vit-D in brain tissue. Here, we present a novel liquid chromatography tandem mass spectrometry (LC-MS/MS) method and for the first time, demonstrate detection of seven major vit-D metabolites in brain tissues of C57BL/6J wild-type mice, namely: 1,25(OH)_2_D_3_, 3-epi-1,25(OH)_2_D_3,_ 1,25(OH)_2_D_2_, 25(OH)D_3_, 25(OH)D_2_, 24,25(OH)_2_D_3_, and 24,25(OH)_2_D_2_. Chromatographic separation was achieved on a pentaflurophenyol column 3 mM ammonium formate with water/methanol [A] and methanol/isopropanol [B] phases. Detection was by positive-ion electrospray tandem mass spectrometry. We used calibration standards of each metabolite prepared in brain matrices to validate the detection range, precision, accuracy and recovery. Isotopically labelled analogues, 1,25(OH)_2_D_3_-d_3_, 25(OH)D_3_-C_5_ and 24,25(OH)_2_D_3_-d_6_, served as the internal standards for the closest molecular related metabolite in all measurements. The calibration range was between 1 fg/mL to 10 ng/mL with an LLOD and LLOQ of 10 fg/mL and 3 fg/mL, respectively. The intra-/inter-day precision and accuracy for measuring brain vit-D metabolites ranged between 0.12-11.53% and 0.28-9.11%, respectively. Recovery ranged between 99.06% and 106.9% for all metabolites. Collectively, the sensitivity and efficiency of our method supersedes previously reported protocols used to measure vit-D and to our knowledge, the first protocol to reveal the abundance of 25(OH)D_2_, 1,25(OH)D_2_ and 24,25(OH)_2_D_2_, in brain tissue of any species. This technique may be important in supporting the future advancement of pre-clinical research into the function of vit-D in neurophysiological, neuropsychiatric, and neurodegeneration.

## 1. Introduction

Increasing evidence indicates that vitamin D (vit-D) is a potent regulator of brain function. A number of epidemiological and experimental studies have implicated vit-D in a wide range of neurological function including, modulating neurotrophic factors, neuronal calcium regulation and signalling, neurotransmission, synaptogenesis, neurogenesis and neuroprotection [1-5]. Moreover, disturbances in cerebral vit-D homeostasis have been linked to neurophysiological and neuropsychiatric disorders such as depression, autism, psychosis, and schizophrenia [6]. However, presently, the homeostasis, metabolism and kinetics of vit-D metabolites in the brain remain largely unknown. Indeed, with techniques that are currently available, only three metabolites of vit-D: 25(OH)D_3_, 24,25(OH)_2_D_3_ and 1,25(OH)_2_D_3_ have been reported in brain tissues to date. Thus, it is important to establish an accurate and reproducible methodologies to quantitate vit-D metabolites in brain.

According to the Vitamin D External Quality Assessment Scheme (DEQAS) comparing methodologies measuring 25(OH)D abundance in biological fluids such as plasma across 871 laboratories in 54 countries, approximately 80% of laboratories used manual or automated immunoassays such as ELISA, whereas 20% used High Performance Liquid Chromatography (HPLC) or Liquid Chromatography with tandem mass spectrometry (LC-MS/MS) [7]. However, DEQAS and several other studies reported high inter-assay and inter-laboratory variability amongst 25(OH)D metabolite immunoassays [8-14]. Immunoassays also require a significantly larger amount of sample in comparison to HPLC or LC-MS/MS, which often may not be suitable for small animal model studies, where sample volume is limited [15]. Furthermore, the possibility of cross-reactivity of the antibody due to the similarity in chemical structure of different vit-D metabolites are potential confounders [16-18]. Immunoassays are generally not able to distinguish between 25(OH)D_2_ or 25(OH)D_3_ due to increased binding affinity to 25(OH)D_3_, reducing the assay detection sensitivities in comparison to HPLC, or LC-MS/MS [7]. The 24,25(OH)2D_3_ metabolite is present in approximately 10–15% of the total 25(OH)D concentration, thus significantly reducing the accuracy of immunoassays [19, 20] and possibly leading to aberrant interpretation of physiological effects.

The use of LC-MS/MS potentially enables the quantitation of various compounds at wide-ranging concentrations with small volumes, mitigating critical issues that may arise with immunodetection methods [21]. Whilst multiple studies have demonstrated mass spectrometry assays for the quantitative measurement of vit-D metabolites in serum [22-25], the analysis of vit-D in lipid-rich brain tissues remains underdeveloped, or not adequately validated. In 2014, Ahonen at el. developed a method utilizing ultra-high-performance liquid chromatography-atmospheric pressure photoionization-tandem mass spectrometry to analyse vit-D related compounds in mouse brain and cell line samples and was the first to report and quantify 25(OH)D_3_ in mouse brain [26]. In 2015 Xue et al. developed a method using high performance liquid chromatography tandem mass spectrometry to simultaneously quantify the concentrations of 25(OH)D_3_ and 24,25(OH)_2_D_3_ in the brain of rats [27]. Fu et al. also developed an ultra-pressure liquid chromatography tandem mass spectrometry and successfully quantified 25(OH)D_3_ and 1,25(OH)_2_D_3_ in porcine brain and human brain [28]. However, the former protocol only reported the presence of 1,25(OH)_2_D_3_ in the prefrontal and middle frontal cortices and did not detect vit-D_3_ in any other regions of the brain. The limitations across these protocols may be due to the use of complex and perhaps incomplete offline extractions, allowing the detection of only 1-2 metabolites at sensitivities in the ng/mL to pg/mL range. Moreover, a C-18 column were used for each of the aforementioned methods, which may not be suitable for complex lipid rich tissue separations such as brain.

In the present study, we developed a highly sensitive and describe a rapid automated online extraction (OLE) method in conjunction with LC-MS/MS coupled to a pentafluorophenyl (PFP). The protocol described herein enabled for the first time, detection and quantification of seven major vit-D metabolites in mouse brain including 1,25(OH)_2_D_3_, 3-epi-1,25(OH)_2_D_3_, 1,25(OH)_2_D_2_, 25(OH)D_3_, 25(OH)D_2_, 24,25(OH) _2_D_3_, and 24,25(OH)_2_D_2_ with markedly greater sensitivity than previously reported protocols (fg/ml).

## 2. Materials and methods

### 2.1. Chemicals

All chemicals used in the protocol were LCMS grade. Acetonitrile (ACN), methanol and water were purchased from Thermofisher (Massachusetts, USA). Ammonium formate (>99%) was purchased from Sigma-Aldrich (Darmstadt, Germany). All standard vit-D compounds and isotope-labelled analogues, 1,25(OH)_2_D_3_, 1,25(OH)_2_D_2_, 24,25(OH)_2_D_3_, 24,25(OH)_2_D_3_-d_6_, 24,25(OH)_2_D_2_, 1,25(OH)_2_D_3_-d_3_, 25(OH)D_3_, 25(OH)D_2_, and 25(OH)D_3_-C_5_ were purchased from PM Separations Pty Ltd (QLD, Australia).

### 2.2. Determination of retention time and collision energies of vit-D precursor and product pairs

Retention time and collision energies were determined via direct infusion of the vit-D standards and isotopes. Each vit-D metabolite, 1,25(OH)_2_D_3_, 1,25(OH)_2_D_2_, 24,25(OH)_2_D_3_, 24,25(OH)_2_D_3_-d_6_, 24,25(OH)_2_D_2_, 1,25(OH)_2_D_3_-d_3_, 25(OH)D_3_, 25(OH)D_2_, and 25(OH)D_3_-C_5_, was prepared by dissolving in ethanol at 1 mg/mL concentration, pipetted into aliquots, and stored in amber glass vials at −80°C. All working solutions were prepared by serial dilution of the stock solutions in ACN. For the determination of retention time and collision energies, standards at 100 ng/mL were used and ran on LC-MS/MS as described in the following section.

### 2.3. LC-MS/MS analysis

Vit-D metabolites were analysed using the EVOQ elite triple quadrupole mass spectrometer with an Advance UPHLC OLE system (Bruker, Billerica, MA) and a CTC HTS-xt autosampler (CTC Analytics AG, Switzerland), where the samples resided in glass vials at 4°C. The mass spectrometer was operated in MS/MS mode with HESI source, capillary voltage set to 3500 volts, cone at 350°C and 20 psi gas, heated probe at 300°C and 50 psi, and nebuliser at 60 psi. Analysis was performed on a poroshell 120 pentafluorophenyl (PFP) column (150 × 2.1 mm, 2.7 µm (Agilent Technologies, Santa Clara, CA, USA) with column oven set to 50°C. Mobile phase A was water with 3 mM ammonium formate, mobile phase B was 2:49:49 water:methanol:isopropanol with 3 mM ammonium formate. The LC was operated at a flow rate of 300 µL/min, with programming shown in table 1.

**Table 1.** Liquid chromatography programming

| Time (min) | Pump B (%) |
| --- | --- |
| 0 | 50 |
| 3 | 50 |
| 15 | 90 |
| 15.1 | 100 |
| 35.1 | 100 |
| 35.2 | 50 |
| 40.2 | 50 |

### 2.4. Online extraction

All standards and samples were run with an online extraction method in order to increase efficiency and throughput whilst simplifying the workflow. Online extraction was performed using a 6-port injection valve and a 10-port switching valve. Flow connections and valve programming are represented in Figure 1. Sample is injected into loop in 1., and loaded onto the trap column in 2. This cycle can be repeated as many times as necessary to concentrate the sample before 3., the analyte is eluted from the trap column onto the analytical column. A poroshell 120 EC-C18 guard column (2.1 × 5 mm, 2.7 mm; Agilent Technologies, Santa Clara, CA, USA) was used to trap vit-D metabolites, before back flushing on to the analytical column. The trap column was equilibrated at 1 mL/min for 1 min with 50% methanol with 3 mM ammonium formate, then injected with 80 μL of prepared brain sample and loaded for 45 s at 1 mL/min. Re-equilibration of the trap column occurred simultaneous to re-equilibration of analytical column, and therefore did not affect the analysis time. This cycle was repeated for a total of 5 trapping cycles resulting in a total volume injection of 400 μL.

**Figure 1.**
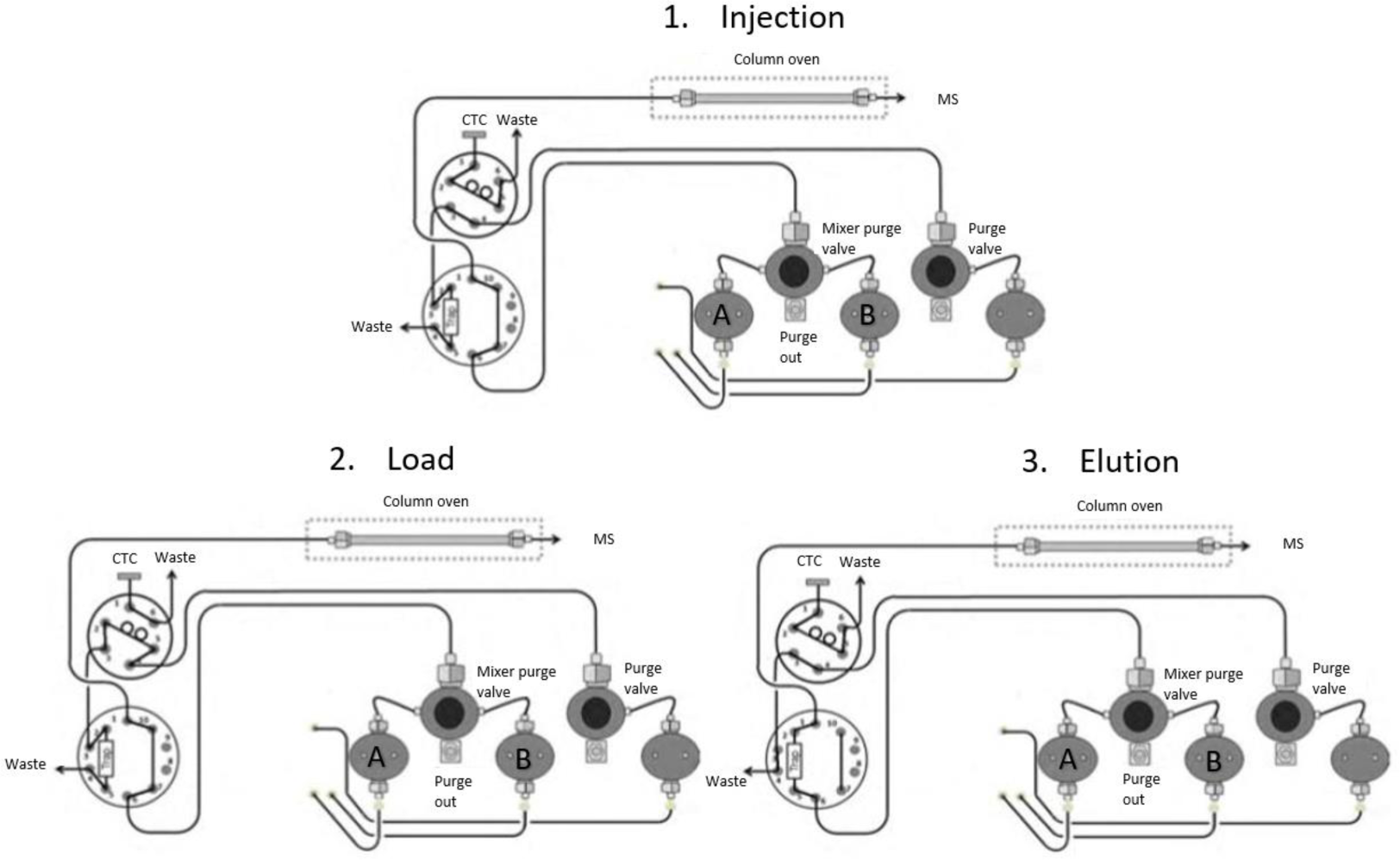
OLE configuration.

### 2.5. Mouse brain sample collection and preparation

Twelve male C57BL/6J mice aged 6 weeks were obtained from the Animal Resource Centre (Murdoch, Western Australia). The animals were maintained in an environmentally controlled animal facility in ventilated cages, temperature 22 °C, 12hr light/dark cycle (Curtin Animal Holding Facility, Curtin University). They were fed standard laboratory chow (AIN93M, Speciality Feeds, Glen Forrest, WA) with water ad libitum. Animals were upheld in accordance with internationally accepted ethical principles for laboratory animal use. All procedures described in this study abided by the Animal Research: Reporting of In Vivo Experiments (ARRIVE) guidelines and were approved by Curtin Animal Ethics Committee (approval no. ARE2018-35).

At 7 weeks of age, the mice were anaesthetized with 2% gaseous isoflurane and euthanised via cervical dislocation [29]. Whole brain specimens were carefully extracted and immediately snap frozen in liquid nitrogen and kept at -80°C until analysis. Whole brain samples were thawed and weighed before being manually ground with ACN using a small pestle. ACN volume was calculated by dividing brain weight (g) by 0.15. Three isotopically labelled standards (25(OH)D_3_-C_5_, 1,25(OH)_2_D_3_-d_3,_ 24,25(OH)_2_D_3_-d_6_) were added for a final concentration of 1.25 ng/mL. The brain homogenate was sonicated in a water bath at 4°C (30 second cycles of 10 s on/20 s off) for 15 min and incubated at 4°C for 10 min to facilitate protein precipitation. The sample was then vigorously vortexed for 5 min at 22°C and centrifuged at 1500 x*g* for 10 min at 4°C. The supernatant from 12 individual mice was then pooled. The sample was stored at -80°C until use. 200 μL of 3 mM ammonium formate in H_2_O was added followed by 1 min vortex immediately prior to analysis.

All following validation analyses were done by spiking vit-D metabolites standards into brain homogenate in order to validate the practicality and applicability of our methods in the mouse brain tissue matrix. The validation analyses were done by subtracting the concentrations of endogenous vit-D metabolites in the blank brain matrix.

### 2.6. Linearity and lower limits of detection/quantification

The calibration curve for each vit-D metabolite was drawn and the LLOD/LLOQ was determined with vit-D standards ranging from 1 fg/mL to 10 ng/mL. The calibration standards were prepared by diluting the stock solution in ACN and added to the brain matrix and ran on LC-MS/MS coupled with online extraction. LLOD was estimated from the linear regression by calculating a concentration equal to triple the averaged non-response with a signal to noise above 30:1. LLOQ was determined by the lowest calibration point with no variation in concentration in the linear regression with a signal to noise above 30:1.

### 2.7. Intra- and inter-day precision and accuracy

The reproducibility of intra- and inter-day measurements were determined with the brain matrix spiked with low, medium, and high standard solution at concentrations of 100, 500 and 1000 pg/mL, respectively. Intraday precision and accuracy were determined with 3 replicate measurements of each low, medium, and high standard. Interday precision and accuracy were determined with repeated measures of low, medium, and high standards over 3 consecutive days. Metabolites were quantified by normalisation to the closest matching internal standard, and subsequent calculation from the constructed matching calibration curves. The same application was used in the quantification of 1,25(OH)_2_D_3_, 1,25(OH)_2_D_2_, 25(OH)D_3_, 25(OH)D_2_ and 24,25(OH)_2_D_3_, and 24,25(OH)_2_D_2_.

Precision was calculated as the total variance of quantified results, while accuracy was determined as the difference between the actual and calculated theoretical concentrations. Precision and accuracy were calculated in both intra- and inter-day periods and expressed as a percentage relative standard deviation (RSD%).

### 2.8. Recovery

Recovery rate of vit-D metabolites during the extraction process was determined by spiking ACN with vit-D standards to a concentration of 100 ng/mL before or after the extraction, then comparing the difference in concentrations. The measurement were repeated in triplicate.

#### Application of the established method to the measurement of endogenous cerebral vit-D metabolites

The concentrations of endogenous brain vit-D metabolites, 1,25(OH)_2_D_3_, 1,25(OH)_2_D_2_, 25(OH)D_3_, 25(OH)D_2_ and 24,25(OH)_2_D_3_, and 24,25(OH)_2_D_2_, were measured in brain tissue homogenates samples spiked with isotopes internal standards. The brain matrix was run on LC-MS/MS with online extraction as described above. The concentration of each metabolite was concurrently determined based on a calibration curve. Seven replicates were run daily, over 3 days.

### 2.9. Statistical analysis

Linearity slope and regression coefficients were determined by linear regression. All statistical analyses were done using Microsoft Excel 2010.

## 3. Results

### 3.1. Determination of retention time and collision energy

The retention time and collision energy of each vit-D metabolite was determined by using vit-D standards. Targeted analytes produced a much stronger signal in positive mode using the electrospray ion source, thus, all the vit-D metabolites were detected under MRM positive mode. The most abundant precursor ions were chosen to increase sensitivity whilst product ions were selected according to greatest intensity and optimal sensitivity. Precursor and product pairs, associated retention times, collision energy and MRM transitions are presented in Table 2 and shown in Figure 2. The OLE-LC-MS/MS method shows good chromatographic performance with all eluting analytes showing clear separation with confident individual detection (Fig. 3).

**Table 2.** Precursor and product pairs and associated retention time and collision energies

| Vitamin D metabolite | Precursor | Products | Retention time (min) | Collision energy (V) | MRM transition (m/z) |
| --- | --- | --- | --- | --- | --- |
| **1,25(OH)<sub>2</sub>D<sub>3</sub>** | 399.30 | 381.2 | 7.02 | 9.0 | 277.8 |
|  |  | 135.1 |  | 21.0 | 277.8 |
| **1,25(OH)<sub>2</sub>D<sub>2</sub>** | 446.30 | 411.3 | 7.36 | 9.0 | 277.8 |
|  |  | 135.1 |  | 11.0 | 277.8 |
| **1,25(OH)<sub>2</sub>D<sub>3</sub>D<sub>3</sub>** | 437.30 | 384.3 | 7.03 | 8.0 | 277.8 |
|  |  | 138.1 |  | 17.0 | 277.8 |
| **24,25(OH)<sub>2</sub>D<sub>3</sub>** | 417.3 | 381.3 | 6.48 | 8.0 | 277.8 |
|  |  | 399.3 |  | 7.0 | 277.8 |
| **24,25(OH)<sub>2</sub>D<sub>2</sub>** | 393.3 | 268.1 | 7.02 | 24.0 | 277.8 |
|  |  | 243.2 |  | 9.0 | 277.8 |
| **24,25(OH)<sub>2</sub>D<sub>3</sub>D<sub>6</sub>** | 423.3 | 405.2 | 6.47 | 9.0 | 277.8 |
|  |  | 387.2 |  | 13.0 | 277.8 |
| **25(OH)D<sub>3</sub>** | 401.30 | 383.3 | 8.98 | 7.0 | 277.8 |
|  |  | 365.3 |  | 8.0 | 277.8 |
| **25(OH)D<sub>2</sub>** | 413.3 | 395.3 | 9.20 | 8.0 | 277.8 |
|  |  | 107.2 |  | 21.0 | 277.8 |
| **25(OH)D<sub>3</sub>-C<sub>5</sub>** | 406.2 | 370.2 | 8.97 | 8.0 | 277.8 |
|  |  | 388.2 |  | 7.0 | 277.8 |

**Figure 2.**
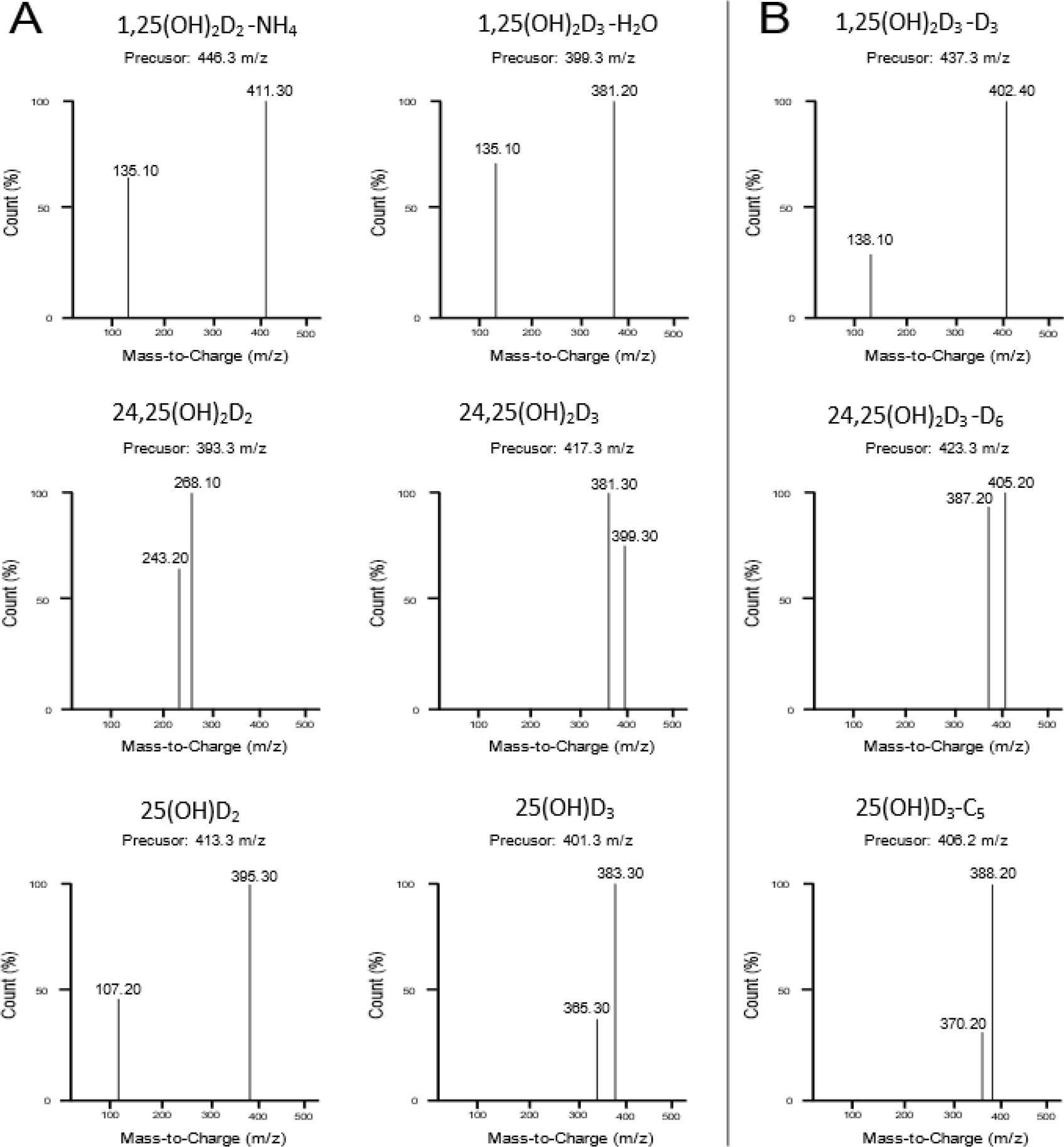
Precursor/product ion spectrum of six vit-D internal standards (A) and associated isotopes (B) in mouse brain

**Figure 3.**
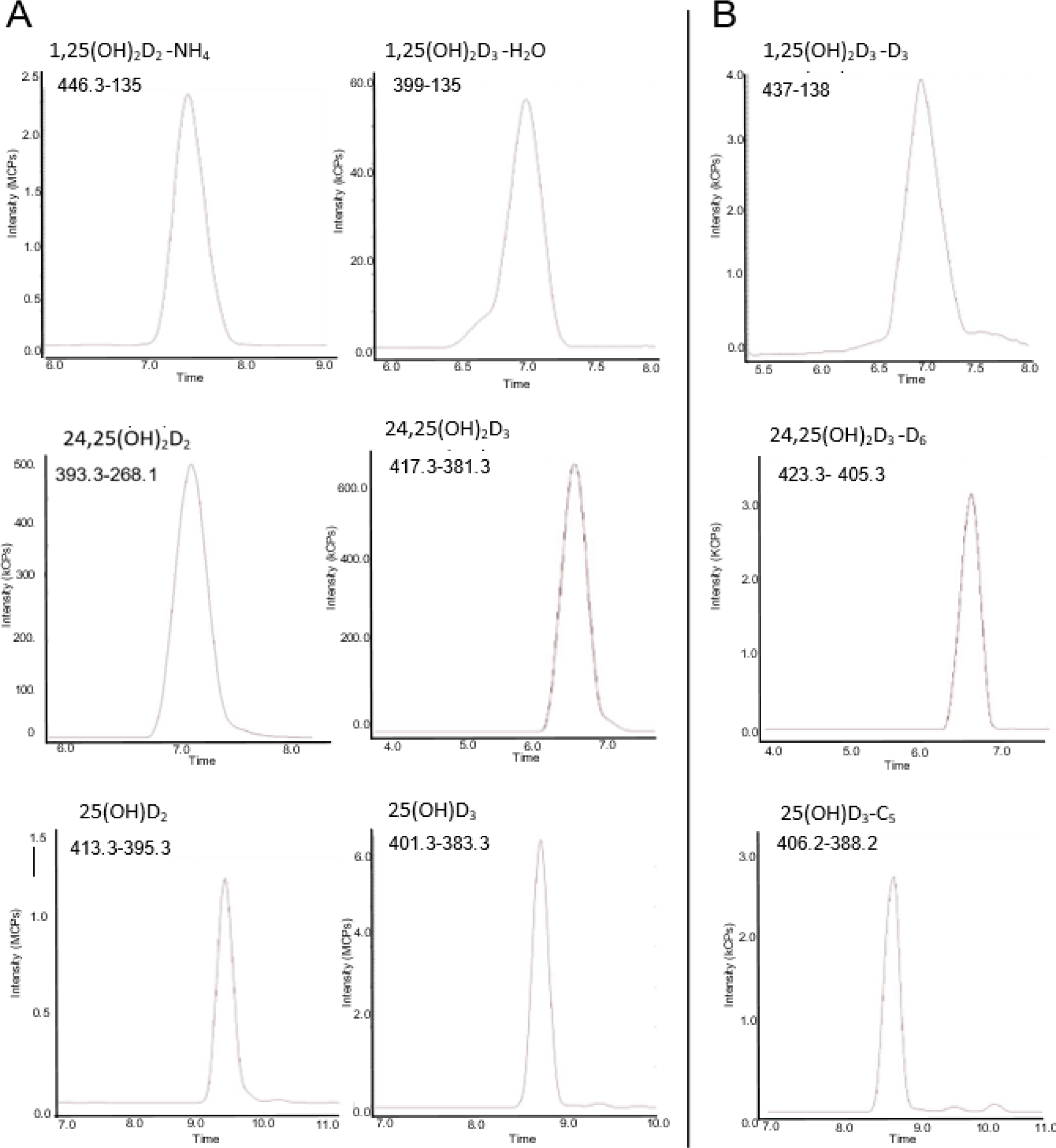
Chromatogram of six vit-D internal standards (A) and associated isotopes (B) in mouse brain at a 100 ng concentration and flow rate of 300 µL/min.

### 3.2. Linearity, limits of detection and quantification

The linearity of the calibration standards ranging from 1 fg/mL to 10 ng/mL, the LLOD, LLOQ and linear range are indicated in Table 3. Linearity R^2^ values within the linear range are also presented in the table. The signal to noise for each metabolite is shown in Table 4. All vit-D standards spiked in the brain matrix were detectable as low as 3 fg/mL and quantifiable as low as 10 fg/mL. Linear range was only assessed as high as 10 ng/mL for all metabolites.

**Table 3.** Calibration linear regression and coefficients of determination (R^2^) for lower limit of detection (LLOD), lower limits of quantification (LLOQ), and linear range of exogenous internal standards and isotopes using a maximum of ten measuring points between from 1 fg/mL to 10 ng/mL

| **Vit-D metabolite** | **R<sup>2</sup>** | **LLOD ( fg/mL)** | **LLOQ ( fg)** | **Linear range (ng)** |
| --- | --- | --- | --- | --- |
| **1,25(OH)<sub>2</sub>D<sub>3</sub>** | 0.9932 | 16.5 | 100 | 10 |
| **1,25(OH)<sub>2</sub>D<sub>2</sub>** | 0.9996 | 16.5 | 100 | 10 |
| **24,25(OH)<sub>2</sub>D<sub>3</sub>** | 0.9989 | 111 | 500 | 10 |
| **24,25(OH)<sub>2</sub>D<sub>2</sub>** | 0.9986 | 111 | 500 | 10 |
| **25(OH)D<sub>3</sub>** | 0.9899 | 3 | 10 | 10 |
| **25(OH)D<sub>2</sub>** | 0.9934 | 111 | 500 | 10 |

**Table 4.**
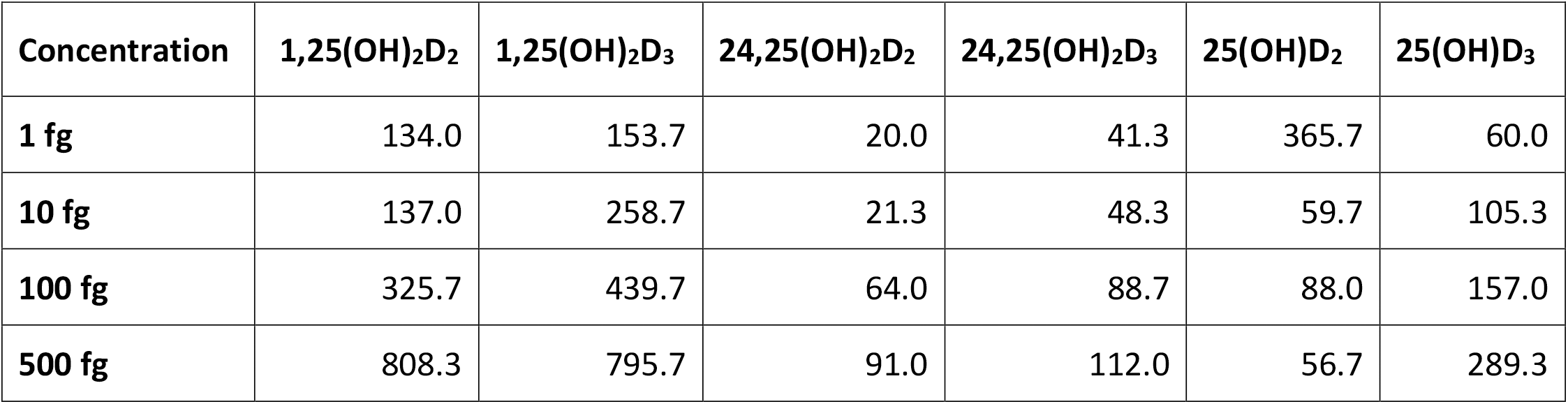
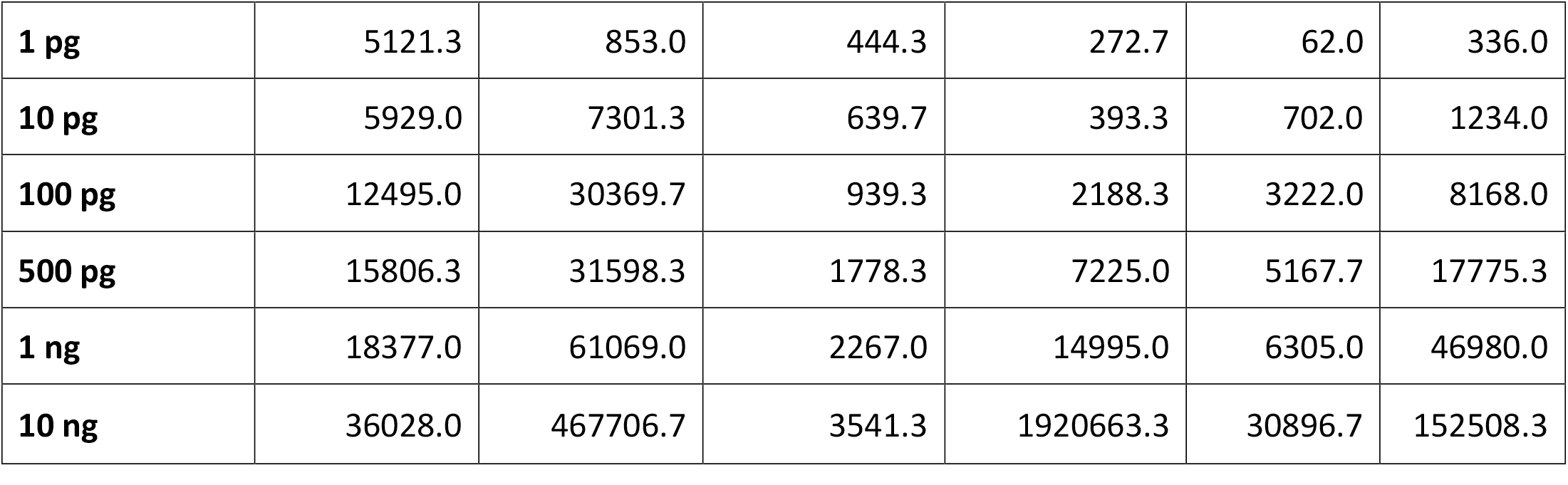
Average signal to noise ratio for six metabolites at nine concentrations taken over the three-day accuracy and precision measures of exogenous brain homogenate

| **Concentration** | **1,25(OH)<sub>2</sub>D<sub>2</sub>** | **1,25(OH)<sub>2</sub>D<sub>3</sub>** | **24,25(OH)<sub>2</sub>D<sub>2</sub>** | **24,25(OH)<sub>2</sub>D<sub>3</sub>** | **25(OH)D<sub>2</sub>** | **25(OH)D<sub>3</sub>** |
| --- | --- | --- | --- | --- | --- | --- |
| **1 fg** | 134.0 | 153.7 | 20.0 | 41.3 | 365.7 | 60.0 |
| **10 fg** | 137.0 | 258.7 | 21.3 | 48.3 | 59.7 | 105.3 |
| **100 fg** | 325.7 | 439.7 | 64.0 | 88.7 | 88.0 | 157.0 |
| **500 fg** | 808.3 | 795.7 | 91.0 | 112.0 | 56.7 | 289.3 |
| **1 pg** | 5121.3 | 853.0 | 444.3 | 272.7 | 62.0 | 336.0 |
| **10 pg** | 5929.0 | 7301.3 | 639.7 | 393.3 | 702.0 | 1234.0 |
| **100 pg** | 12495.0 | 30369.7 | 939.3 | 2188.3 | 3222.0 | 8168.0 |
| **500 pg** | 15806.3 | 31598.3 | 1778.3 | 7225.0 | 5167.7 | 17775.3 |
| **1 ng** | 18377.0 | 61069.0 | 2267.0 | 14995.0 | 6305.0 | 46980.0 |
| **10 ng** | 36028.0 | 467706.7 | 3541.3 | 1920663.3 | 30896.7 | 152508.3 |

### 3.3. Intra- and inter-day repeatability and recovery

Intra-day precision and accuracy (RSD%) for vit-D metabolite standards in brain matrix is presented in Table 5. The precision and accuracy ranged from 80 to 120%. The precision in the inter-day measures for all metabolites were between 12.47% and –9.53% (Table 6). The recovery ranged between 99.06% and 106.9% (Table 7).

**Table 5.**
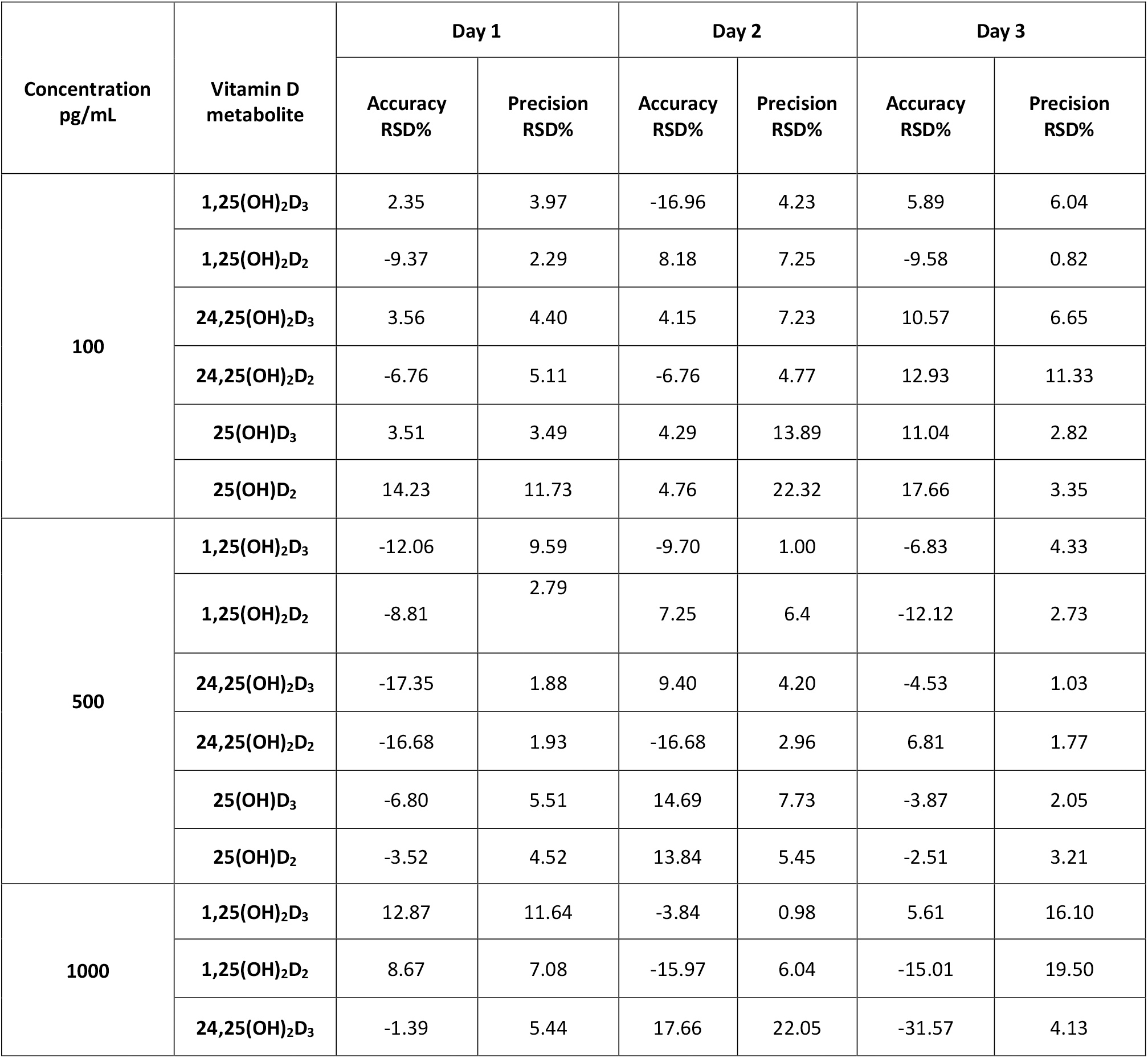
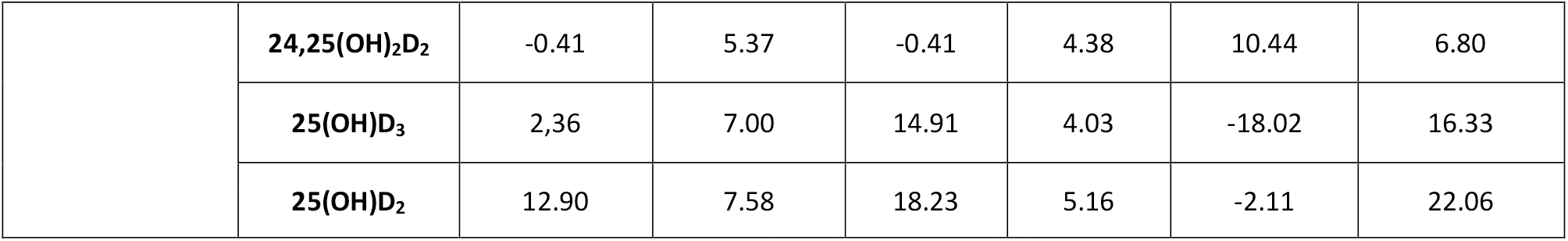
Intraday accuracy and precision (RSD %)

**Table 6.** Inter-day accuracy and precision (RSD %)

| Concentration<br>pg/mL | Vitamin D<br>metabolite | Inter-day |  |
| --- | --- | --- | --- |
|  |  | Accuracy<br>RSD % | Precision<br>RSD % |
| **100** | 1,25(OH)<sub>2</sub>D<sub>3</sub> | -2.90 | 4.75 |
|  | 1,25(OH)<sub>2</sub>D<sub>2</sub> | -3.59 | 3.45 |
|  | 24,25(OH)<sub>2</sub>D<sub>3</sub> | 6.10 | 6.09 |
|  | 24,25(OH)<sub>2</sub>D<sub>2</sub> | -0.20 | 7.07 |
|  | 25(OH)D<sub>3</sub> | 6.28 | 6.73 |
|  | 25(OH)D<sub>2</sub> | 12.22 | 12.47 |
| **500** | 1,25(OH)<sub>2</sub>D<sub>3</sub> | -9.53 | 4.97 |
|  | 1,25(OH)<sub>2</sub>D<sub>2</sub> | -4.56 | 3.98 |
|  | 24,25(OH)<sub>2</sub>D<sub>3</sub> | -4.16 | 2.37 |
|  | 24,25(OH)<sub>2</sub>D<sub>2</sub> | -8.85 | 2.22 |
|  | 25(OH)D<sub>3</sub> | 1.34 | 5.10 |
|  | 25(OH)D<sub>2</sub> | 2.60 | 4.47 |
| **1000** | 1,25(OH)<sub>2</sub>D<sub>3</sub> | 4.88 | 9.58 |
|  | 1,25(OH)<sub>2</sub>D<sub>2</sub> | -7.44 | 10.88 |
|  | 24,25(OH)<sub>2</sub>D<sub>3</sub> | -5.10 | 10.54 |
|  | 24,25(OH)<sub>2</sub>D<sub>2</sub> | -3.0 | 5.52 |
|  | 25(OH)D<sub>3</sub> | -0.25 | 9.12 |
|  | 25(OH)D<sub>2</sub> | 4.47 | 11.60 |

**Table 7.** Recovery (%) of metabolites showing mean, SD and RSD (%)

|  |  | 1,25(OH)<sub>2</sub>D<sub>2</sub> | 1,25(OH)<sub>2</sub>D<sub>3</sub> | 24,25(OH)<sub>2</sub>D<sub>3</sub> | 24,25(OH)<sub>2</sub>D<sub>2</sub> | 25(OH)D<sub>2</sub> | 25(OH)D<sub>3</sub> |
| --- | --- | --- | --- | --- | --- | --- | --- |
| Prespike | Mean | 5.32 | 0.15 | 2.17 | 1.73 | 1.32 | 11.89 |
|  | SD | 0.31 | 0.00 | 0.08 | 0.05 | 0.04 | 0.50 |
|  | RSD % | 5.75 | 2.20 | 3.53 | 3.15 | 3.34 | 4.18 |
| Postspike | Mean | 5.76 | 0.15 | 2.32 | 1.71 | 1.36 | 12.71 |
|  | SD | 5.76 | 0.15 | 2.32 | 1.71 | 1.36 | 12.71 |
|  | RSD % | 2.98 | 0.08 | 1.44 | 0.98 | 0.65 | 8.05 |
|  | Recovery % | 108.32 | 105.39 | 106.60 | 99.09 | 103.01 | 106.92 |

### 3.4. Measurement of endogenous brain vit-D metabolites

Mean metabolite concentrations, standard deviation and intra- and inter-day precision (RSD%) measures for endogenous vit-D are presented in Table 8. Intra- and inter-day precision for all metabolites ranged between 0.12-11.53% and 0.28-9.11%, respectively.

**Table 8.** Mean, standard deviation and RSD % measures of endogenous homogenate (pg/g)

|  |  | 1,25(OH)<sub>2</sub>D<sub>2</sub> | 1,25(OH)<sub>2</sub>D<sub>3</sub> | 24,25(OH)<sub>2</sub>D<sub>2</sub> | 24,25(OH)<sub>2</sub>D<sub>3</sub> | 25(OH)D<sub>2</sub> | 25(OH)D<sub>3</sub> |
| --- | --- | --- | --- | --- | --- | --- | --- |
| Intraday |  |  |  |  |  |  |  |
| Day 1 | Mean pg/g | 24.99 | ND | 234.77 | 91.54 | 127.93 | 19.72 |
|  | SD | 0.42 |  | 15.32 | 0.11 | 0.80 | 0.34 |
|  | % RSD | 1.68 |  | 6.53 | 0.12 | 0.62 | 1.70 |
| Day 2 | Mean pg/g | 25.19 | ND | 205.48 | 92.05 | 114.85 | 18.43 |
|  | SD | 0.42 |  | 23.69 | 0.22 | 0.87 | 0.31 |
|  | % RSD | 1.67 |  | 11.53 | 0.23 | 0.76 | 1.66 |
| Day 3 | Mean pg/g | 24.49 | ND | 222.66 | 92.78 | 143.41 | 13.77 |
|  | SD | 0.23 | ND | 20.65 | 0.44 | 4.43 | 1.41 |
|  | % RSD | 0.94 | ND | 9.27 | 0.48 | 3.09 | 10.21 |

| Interday |  |  |  |  |  |  |  |
| --- | --- | --- | --- | --- | --- | --- | --- |
| Average | Mean<br>pg/g | 24.89 | ND | 220.97 | 92.12 | 128.73 | 17.31 |
|  | SD | 0.36 |  | 19.89 | 0.26 | 2.03 | 0.68 |
|  | %<br>RSD | 1.43 |  | 9.11 | 0.28 | 1.49 | 4.53 |

\*ND= Not detected

Five vit-D metabolites were detected endogenously within the validated detection range. These include 1,25(OH)_2_D_2_, 25(OH)D_3_, 25(OH)D_2_ and 24,25(OH)_2_D_3_, and 24,25(OH)_2_D_2_. Whilst 1,25(OH)_2_D_3_ was not detected at the validated retention time, an unknown peak was present in the chromatogram at 7.5 min as shown in Fig 4. We confirmed this peak to be the epimer of 1,25(OH)_2_D_3_ (3-epi-1,25(OH)_2_D_3_) as described below in Section 3.4.1.

**Figure 4.**
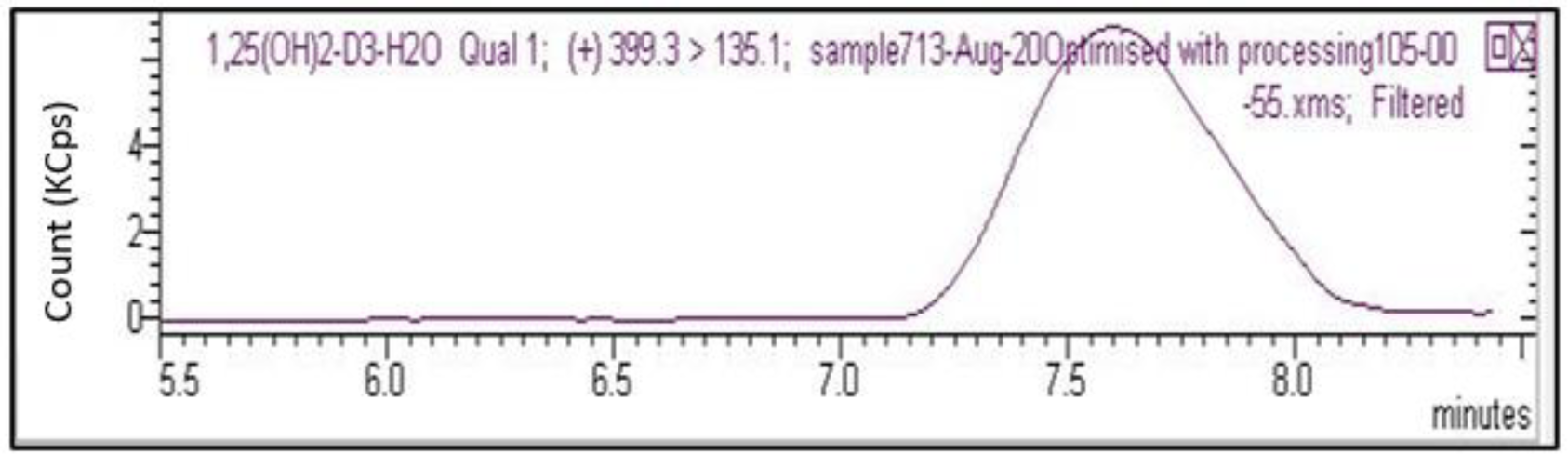
An unknown 7.5 min peak was observed within the 1,25(OH)_2_D_3_ chromatogram for all seven endogenous samples. This was later validated as 3-epi-1,25(OH)_2_D_3_.

#### 3.4.1. Detection and validation of 3-epi-1,25(OH)2D3 metabolite

We identified an unknown, but consistent peak at 7.5 min in the 1,25(OH)_2_D_3_ chromatogram (Fig 4). We validated this as the epimer of 1,25(OH)_2_D_3_ as per the protocol described in Section 2.2, using a 3-epi-1,25(OH)_2_D_3_ standard (abcr, Germany cas 61476-45-7).

3-epi-1,25(OH)_2_D_3_ precursor and product pairs and collision energy are listed in table 9. To validate the retention time, spiked known concentrations (1.25 ng) of 3-epi-1,25(OH)_2_D_3_ standard were run in both ACN and brain matrix, and brain matrix calibration (1.6-100 ng) confirming the 7.5 min retention time (table 10).

**Table 9.** 3-epi-1,25(OH)_2_D_3_ precursor and product, retention time and collision energy

| Precursor | Products | Retention time<br>(min) | Collison<br>energy (V) | MRM<br>transition<br>(m/z) |
| --- | --- | --- | --- | --- |
| 399.20 | 199.10 | 7.5 | 13.0 | 277.8 |
|  | 299.10 |  | 9.0 | 277.8 |

**Table 10.**
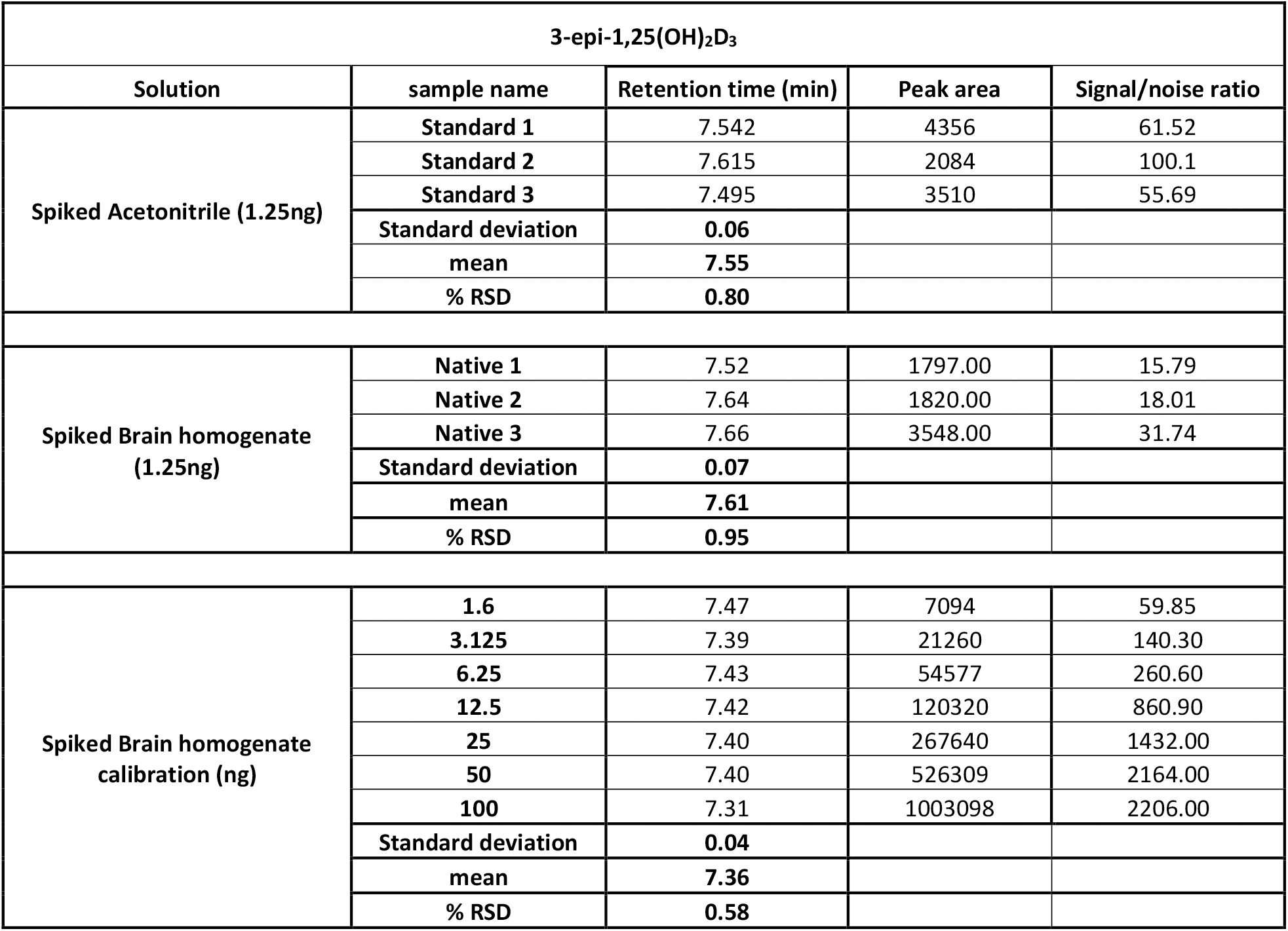
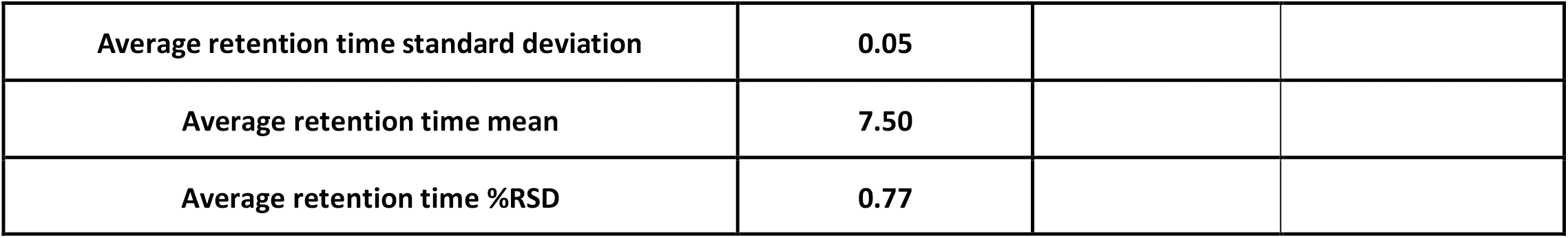
3-epi-1,25(OH)_2_D_3_ retention time validation.

| 3-epi-1,25(OH)<sub>2</sub>D<sub>3</sub> |  |  |  |  |
| --- | --- | --- | --- | --- |
| Solution | sample name | Retention time (min) | Peak area | Signal/noise ratio |
| Spiked Acetonitrile (1.25ng) | Standard 1 | 7.542 | 4356 | 61.52 |
|  | Standard 2 | 7.615 | 2084 | 100.1 |
|  | Standard 3 | 7.495 | 3510 | 55.69 |
|  | Standard deviation | 0.06 |  |  |
|  | mean | 7.55 |  |  |
|  | % RSD | 0.80 |  |  |
| Spiked Brain homogenate<br>(1.25ng) | Native 1 | 7.52 | 1797.00 | 15.79 |
|  | Native 2 | 7.64 | 1820.00 | 18.01 |
|  | Native 3 | 7.66 | 3548.00 | 31.74 |
|  | Standard deviation | 0.07 |  |  |
|  | mean | 7.61 |  |  |
|  | % RSD | 0.95 |  |  |
| Spiked Brain homogenate<br>calibration (ng) | 1.6 | 7.47 | 7094 | 59.85 |
|  | 3.125 | 7.39 | 21260 | 140.30 |
|  | 6.25 | 7.43 | 54577 | 260.60 |
|  | 12.5 | 7.42 | 120320 | 860.90 |
|  | 25 | 7.40 | 267640 | 1432.00 |
|  | 50 | 7.40 | 526309 | 2164.00 |
|  | 100 | 7.31 | 1003098 | 2206.00 |
|  | Standard deviation | 0.04 |  |  |
|  | mean | 7.36 |  |  |
|  | % RSD | 0.58 |  |  |

|  |  |
| --- | --- |
| **Average retention time standard deviation** | **0.05** |
| **Average retention time mean** | **7.50** |
| **Average retention time %RSD** | **0.77** |

Mean concentrations, standard deviation and intra- and inter-day precision (RSD%) measures for validated endogenous 3-epi-1,25(OH)2D3 are presented in Table 11. Intra-day precision for 3-epi-1,25(OH)_2_D_3_ ranged between 2.53-8.53% and inter-day precision was 6.45%.

**Table 11.** Mean, standard deviation and RSD % measures of endogenous 3-epi-1,25(OH)_2_D_3_ homogenate (pg/g)

| 3-epi-1,25(OH)<sub>2</sub>D<sub>3</sub> |  |  |  |
| --- | --- | --- | --- |
| Intraday |  |  |  |
|  | Mean (pg/g) | SD | RSD % |
| **Day 1** | 227.09 | 19.38 | 8.53 |
| **Day 2** | 228.21 | 18.92 | 8.29 |
| **Day 3** | 268.27 | 6.79 | 2.53 |
| Interday |  |  |  |
|  | Mean (pg/g) | SD | RSD % |
| **Average** | 241.19 | 15.03 | 6.45 |

## 4. Discussion

This study describes a rapid, sensitive OLE-LC-MS/MS protocol on PFP columns that separates and quantifies seven vit-D metabolites 1,25(OH)_2_D_3_; 3-epi-1,25(OH)_2_D_3;_ 1,25(OH)_2_D_2_; 25(OH)D_3_; 25(OH)D_2_; 24,25(OH)_2_D_3_ and 24,25(OH)_2_D_2_ in mouse brain.

Previous studies reporting vit-D metabolites in brain have used offline extraction techniques requiring extensive overnight derivatization, solid-phase extraction, and liquid–liquid extraction, to quantify vit D metabolites with LCMS methods [26-28, 30]. We extend on those studies and describe herein a simplified, more sensitive and robust online extraction process, enabling a stacked loading protocol to effectively concentrate the abundance of quantification of vit D metabolites in brain. The protocol detailed addresses potential significant confounders, and achieves markedly improved sensitivity and identification of brain vit D metabolites that may be physiologically important.

C-18 columns have been historically used in LCMS protocols to determine vit D metabolites ([26-28]. The C18 matrix has preferential retention of hydrophobic moieties within the stationary phase, notionally enabling detection of vit-D metabolites[31]. However, commonly residual abundance of brain phospholipids from offline extracts with direct loading thereafter to C18 columns may markedly interfere with target vit-D metabolite detection and/or significantly increase the threshold detection requirements [32].

The online extraction component of the novel OLE-LC-MS/MS method described with selective elution from the trap column, enables effective bulk decontamination of phospholipids[33]. With the option to loop load and stack, effectively concentrating metabolites of interest, coupled with phospholipid removal, sensitivity appears to be improved orders of magnitude and other metabolites of vit-D with potentially important biological function are realised. Indeed, we report new measures of less-polar metabolites of vit D in mouse brains using the OLE-LCMS on PFP. We suggest that the rigidity of the PFP endcapping provides better shape selectivity, cation exchange, charge transfer, electrostatic interactions, hydrogen bonding, dipole and pi-pi interactions, which allows for greater separation of structural isomers of vit-D compared to C18 reverse phase methods[34].

In the protocol detailed, solvents were selected to utilise the greater hydrogen bonding capabilities of the PFP column compared to C18 columns. We report that water:methanol:isopropanol (2:49:49) with 3 mM ammonium formate obtained the highest recovery for all 7 vit-D metabolites reported. Different concentrations of ammonium formate (3 mM, 6 mM and 10 mM) were tested during the method development processes and 3 mM was chosen for the maximum yield. The method described had excellent limits of detection and quantification with a range of 3-111 fg/mL^-1^ and 10-500 fg/mL^-1,^ respectively. Furthermore, recovery rate of all metabolites ranged between 99.06% and 106.9%. Collectively, this supports the selectivity and sensitivity of the protocol described.

The method detailed here demonstrates a high rate of accuracy and is congruent with previous methods being well within the general range of 80-120% [26]. Considering our wide detection range in complex brain matrix, the results indicate appropriate accuracy of the method by means of repeatability. Furthermore, our targeted vit-D cerebral metabolic profiling method removes any unwarranted sample extraction procedure providing unbiased and robust results, with minimum sample preparation.

## 5. Conclusion

The critical role of vit-D in central nervous system function and evidence of an association with some neurological disorders and neurodegenerative diseases is increasingly indicated. Indeed, the active form of vit-D, 1,25(OH)D_3_ and key enzymes for vit-D metabolism such as CYP27B1 have been identified in cells of the central nervous system. However, only three vit-D metabolites have been reported in brain tissue specimens (25(OH)D_3_, 24,25(OH)_2_D_3_ and 1,25(OH)D_3_) to date. Clearly, more robust and sensitive methods than are currently available are required to understand cerebral vit D homeostasis and if this associated with brain function.

This study validates a relatively simple, sensitive, rapid and specific OLE-LC-MS/MS method to screen and quantify vit-D brain metabolites in brain of C57BL/6J mice.

## Acknowledgements

This work was supported by the National Health and Medical Research Council of Australia; MSWA (Multiple Sclerosis, Western Australia) and the McCusker Charitable Research Foundation.

## Contributions to the field statement

Current and emerging data collected from experimental, pre-clinical and clinical studies have demonstrated disturbances in serum vit-D homeostasis as a major risk in the development and progression neurodegenerative disorders. Whilst the detrimental impact of vit-D deficiency throughout the peripheral and central nervous system is well established, little is known regarding the impact of vit-D dyshomeostasis within the brain. Presently, the reference range for vit-D serum levels are set to maintain skeletal health, while an appropriate concentration for vit-D within the central nervous system remain to be elucidated. Classification of what vit-D levels are recommended are ambiguous and controversial, whilst any mechanisms for potential adverse effects of exaggerated vit-D levels on CNS function remain to be investigated. This protocol significantly improves currently available method of cerebral vit-D detection, developed to the use of OLE-LC-MS/MS method with PFP column. This expanded the detectable range of vit-D metabolites to fg/mL range, which is ∼1,000-fold improvement. It demonstrates cerebral detection of 7 vit D metabolites, as opposed to two previously reported. This is critical in allowing researchers to explore both the fundamental role of vit-D synthesis, metabolism, and homeostasis within the brain and furthering our understanding into neurological and neurodegenerative pathologies.

